# Effect of SNEDDS Loading Pentagamavunon-0 on Memory Impairment and Neurogenesis in Mice with Monosodium Glutamate-Induced Alzheimer’s Disease-Like Symptoms

**DOI:** 10.1101/2024.05.17.594779

**Authors:** Yance Anas, Ratna Asmah Susidarti, Ronny Martien, Nunung Yuniarti

## Abstract

Pentagamavunon-0 (PGV-0), a curcumin analog, has antioxidant and HDAC2 inhibitory properties. It affects neurogenesis and cognitive function in Alzheimer’s disease (AD). Targeting neurogenesis is a strategy currently being developed for AD treatment. This study investigates the therapeutic potential of self-nanoemulsifying drug delivery systems loaded with PGV-0 (SNEDDS PGV-0) on spatial learning, memory impairment, and gene expression-related neurogenesis in mice with monosodium glutamate (MSG) induced AD-like symptoms. MSG 4 g/kg was injected (sc.) into 4-week-old Balb/C mice seven times every alternate day, followed by treatment for 60 days with CMC-Na 0.5%, PGV-0 suspension, SNEDDS PGV-0, and donepezil-HCl. The open-field test, novel object recognition, and 8-radial arm maze test were employed to assess the neurobehavioral performance of mice. Neurogenesis-related gene expression (dcx, nestin, Hes5, and NFIA) was quantified through qPCR analysis. Spatial learning and memory in mice declined with MSG treatment. SNEDDS PGV-0 effectively alleviates mice’s cognitive and memory deficits, as shown in improvements in behavioral parameters, including the Discrimination Index, Recognition Index, and Memory Score. It also restores mRNA expression of several essential neurogenesis-related genes, including dcx, Hes5, and NFIA. SNEDDS PGV-0 demonstrates promising potential as a neurogenesis promoter, making it a viable drug candidate for AD or other neurodegenerative pathologies.

## 1. INTRODUCTION

Alzheimer’s disease (AD) is a severe neurodegenerative disorder that significantly disrupts mental health and cognitive function [1–3]. Amyloid-β (Aβ) plaques and neurofibrillary tangles, two hallmarks of AD, can trigger degeneration and neuronal cell death [4,5]. Beyond these markers, several factors, such as neuroinflammation, oxidative stress, and cholinergic damage, intricately contribute to the persistent progression of neurodegeneration [6,7]. Early neuronal cell death is recognized from the preclinical phase to the late stages of dementia [8]. Therefore, interventions targeting neuroprotection and neurogenesis have emerged as essential strategies to restore cognitive function in AD [9–11]. Recent studies have highlighted a significant decline in neurogenesis among AD patients, particularly in the hippocampus, a critical region for memory formation [12]. Instead, individuals who do not have neurological issues display strong neurogenesis in that same region [13]. It is promising that studies in mice with elevated adult hippocampal neurogenesis have shown positive results in improving cognitive functions such as object recognition, spatial learning, contextual fear conditioning, and extinction learning [9]. These findings suggest that revitalizing adult hippocampal neurogenesis holds enormous potential as an effective therapeutic target for AD. The various pharmacological activities of pentagamavunon-0 (PGV-0), an analog of curcumin, have been described in several studies [14–19], including its role as an antioxidant. Its chemical structure includes phenolic hydroxy and methoxy groups in the ortho position of both aromatic rings, significantly contributing to its potent antioxidant properties and free radical scavenging abilities [20,21]. This antioxidant effect offers the possibility of developing PGV-0 as a potential agent for neurodegenerative disease treatment, including AD [22]. PGV-0 appears to mitigate neuronal damage and prevent neuronal cell death induced by oxidative stress; a phenomenon triggered by the accumulation of Aβ around neurons [23,24]. In addition, PGV-0 exhibits promising potential as an anti-Alzheimer’s drug candidate due to its role as a histone deacetylase-2 (HDAC2) inhibitor. In vitro studies have shown that PGV-0 was 181 times more potent than curcumin in inhibiting HDAC2 activity [25].

The HDAC2 plays a crucial role in the impairment of cognitive function and deficits in adult neurogenesis observed in AD [26–28]. Research has shown that increased levels of HDAC2 expression are associated with decreased short-term memory and synaptic plasticity [29]. Encouragingly, studies suggest that HDAC inhibitors effectively address dysregulation in histone acetylation in Alzheimer’s mouse models. These inhibitors enhance neurogenesis, dendrite growth, synaptic density, and memory formation [30]. In addition, they mitigate Tau protein hyperphosphorylation and reduce neuroinflammation. Selective HDAC2 inhibitors show remarkable potential in promoting dendrite growth and aiding in the degradation of Aβ aggregates. Specific HDAC inhibitors, such as valproic acid, 4-phenylbutyrate, trichostatin A, and suberoylanilide hydroxamic acid, have shown the ability to restore histone acetylation, enhance memory formation, and relieve AD symptoms in animal models. In particular, suberoylanilide hydroxamic acid successfully reversed memory decline, while 4-phenylbutyrate reduced Tau protein hyperphosphorylation and restored acetylation levels of histone H3 and H4 in mouse models of AD [31].

PGV-0 has been classified as a BCS Class II compound because it has low solubility in water [32] and high permeability [33], resulting in limitations on bioavailability owing to degradation and first-pass metabolism [34]. To improve the bioavailability of PGV-0, a lipid-based nanocarrier approach, such as SNEDDS, was employed [35]. In a study, PGV-0 nanoemulsion treatment successfully suppressed the expression of the HDA2 enzyme in ethanol-induced brain disorder in mice [36]. These findings suggest that incorporating PGV-0 in SNEDDS enhances its bioavailability and facilitates its distribution in the brain, allowing for precise targeting of its intended action. However, the specific contributions of SNEDDS PGV-0 to neurogenesis mechanisms in AD animal models, specifically through HDAC2 inhibition, remain to be explored. This study aims to evaluate SNEDDS PGV-0’s effect on improving learning and memory deficits in mice with MSG-induced AD-like symptoms. In addition, we will analyze brain samples to assess PGV-0 impact on neurogenesis-related genes such as dcx, nestin, Hes5, and NFIA.

## 2. MATERIAL AND METHOD

### Material

Commercial monosodium glutamate (PT. Ajinomoto Tbk. Indonesia), vanillin (Merck, Germany), cyclopentanone (Merck, Germany), chloride acid 37% (Merck, Germany), ethanol (Merck, Germany), distilled water (PT. Brataco chemika, Indonesia), mygliol, tween 80, PEG 400 (CV. Chlorogreen Gemilang in Bandung, Indonesia), CMC-Na (Planet Kimia, Indonesia), Donepezil-HCl (PT. Dankos Farma Tbk. Indonesia), phosphate buffer saline (PT. Nitra Kimia, Yogyakarta, Indonesia), FavorPrep™ RNA stabilization solution (Favorgen Biotech corp., Taipei, Taiwan), FavorPrep™ Tissue Total RNA Mini Kit (Favorgen Biotech corp., Taipei, Taiwan), ExcelRT™ Reverse Transcription Kit 100RX (Smobio Technology Inc.), ExcelTaq™ 2x PCR Master Dye Mix 100 RXN (Smobio Technology Inc.).

### 2.2 Animal and handling

Balb/C mice, four-week-old, body weight: 15-25 g, were used as Rodden model and housed in plastic cages measuring 30 cm x 20 cm x 20 cm, with five male and five female mice in each group. The mice were kept according to the animal research guidelines of the Faculty of Pharmacy at Universitas Gadjah Mada, Indonesia. The environmental conditions were carefully maintained at a temperature of 25±2°C, humidity of 55%, and a 12/12-hour dark/light cycle. Comfeed BR1^®^ pellets (PT. JAPFA Comfeed Indonesia Tbk.), containing 4.5% dietary fiber, 21.0% protein, and 4.0% fat, were used to feed the mice. In addition, tap water was provided to the mice to drink during the study, and their weight was measured every five days.

### 2.3 Preparation of monosodium glutamate (MSG) 240 mg/ml

A stock solution of MSG at a concentration of 240 mg/ml was prepared by dissolving 12 g of MSG in 50.0 ml of distilled water.

### 2.4 Synthesis of PGV-0

PGV-0 was synthesized with vanillin and cyclopentanone as starting materials, following a protocol based on an improved technique developed by Ritmaleni [37]. The modification entailed a short reaction (2 hours), followed by washing the crude PGV-0 with distilled water and ethanol. Next, the compound was recrystallized using a two-solvent approach using ethanol and cold-distilled water.

### 2.5 Preparation of PGV-0 loaded SNEDDS

A self-nanoemulsifying drug delivery system containing PGV-0 (SNEDDS PGV-0) was formulated at a concentration of 8.5 mg/ml using mygliol, Tween 80, and PEG 400 as excipients, resulting in a final volume of 10 ml. The preparation involved mixing 85 mg of PGV-0 with 1.50 ml of mygliol and vortexing for 60 seconds. Next, we added 7.5 ml of Tween 80 and 1.0 ml of PEG 400 to the mixture and vortexed it for 60 seconds. Afterward, we sonicated the resulting blend for 20 minutes until it was clear. Before administration, the SNEDDS PGV-0 was dispersed in aquabidest and sonicated for two minutes to achieve final concentrations of 4.25 mg/ml and 2.125 mg/ml.

### 2.6 Preparation of PGV-0 suspension

A suspension of PGV-0 (2.125 mg/ml) was prepared by suspending 21.25 mg of PGV-0 in 10.0 ml of 0.5% CMC-Na solution. The mixture was stirred until homogeneous. Before administration to the mice, the PGV-0 suspension was gently re-suspended.

### 2.7 Preparation of donepezil-HCl 0.18 mg/ml

The mice were administered a daily dose of 3 mg/kg of Donepezil-HCl. This solution was prepared as a stock solution of 0.18 mg/ml. In brief, 1.80 mg of Donepezil-HCl was carefully weighed and dissolved in 10.0 ml of distilled water. The solution was freshly prepared daily.

### 2.8 Design of experimental

Sixty-four-week-old Balb/C mice were allocated into six groups, each consisting of five males and five females housed in separate cages. Groups II to VI received subcutaneous injections of MSG (4 g/kg) every other day for seven doses. The healthy control (HC) group (Group I), received subcutaneous injections of 0.5 ml/30 g of aquabidest every alternate day for seven doses. During the 60-day treatment period, mice in groups II to VI were treated daily by oral gavage with CMC-Na (0.5 ml/30 g), SNEDDS PGV-0 (20 and 40 mg/kg), PGV-0 suspension (40 mg/kg), and Donepezil-HCl (3 mg/kg). During weeks seven and eight of the study, behavioral tests were conducted applying the open-field test (OFT), novel object recognition task (NORT), and 8-radial arm maze (8-RAM) test to measure locomotors activity, anxiety-like behavior, and learning/memory function in mice. Following the final dose on day 60, mouse brains were isolated and stored in an RNA stabilization solution at -20°C for further analysis. The expression levels of the dcx, nestin, Hes5, and NFIA genes, which are associated with neurogenesis, respectively, were measured through quantitative polymerase chain reaction (qPCR).

### 2.9 Open-field test (OFT)

During the experiment, we employed a black, square open box, 38 x 30 x 30 cm, as an open-field arena. The box had no bedding, and we placed the mice in the right-hand corner, facing the wall, and allowed them to explore for five minutes before returning them to their home cages. A camera (Kogan^®^ 4K Action Cam) was used to record the total distance traveled, immobility time, rearing, and grooming time of the mice [38], with video analysis performed using the open-source software Tracker^®^ [39]. After each observation, we cleaned the arena with 70% ethanol and dried with a tissue to remove olfactory signals.

### 2.10 Novel object recognition task (NORT)

Our study employed the NORT procedure, composed of three phases: habituation, familiarization, and testing, as described in Singh *et al*. [40]. To record the behavior of the mice, we used a black open box (38 × 30 × 30 cm) without the ability to view the outside from the inside. A camera was positioned 40 cm above the center of the box. We used two objects to stimulate the mice’s interest: a brown cylinder, 8 x 4 cm, referred to as the familiar object, and a 6 x 8 cm doll of three different colors, the novel object. The experiment started with a 10-minute familiarization phase in an empty arena. After a 20-minute break, the mice were reintroduced to the arena with two familiar objects. We observed the mice for 10 minutes, which represented the familiarization phase. Two hours later, we replaced the familiar object with a novel object. The mice were placed between the objects and observed for five minutes (test phase). We used a video tracking system (Tracker^®^) to record the mice’s interaction time (exploring, touching, and sniffing) with both objects. This data was then utilized to calculate the Discrimination Index (DI) and the Recognition Index (RI) as described in equations (1) and (2) [41].

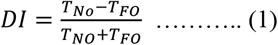

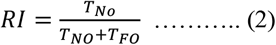

T_NO_: mice’s interaction with novel object

T_FO_: mice’s interaction with familiar object

### 2.11 8-Radial arm maze (8-RAM) test

Our study used an 8-RAM apparatus made of stainless steel to assess mice’s spatial learning and memory. The maze had eight horizontal arms, each measuring 50 cm in length, 10 cm in width, and 10 cm in height. There was an entrance door, a reward cave at the end of each arm, and a circular central zone that connected all the arms with a diameter of 30 cm. The camera was positioned 250 cm high, and the maze was 100 cm above the ground. The experiment was divided into three phases: two days of habituation, four days of training trials, and one day of task testing [40]. The apparatus was disinfected with 70% ethanol and dried with a tissue at the end of each session. The mice were fasted for 24 hours before each session to increase motivation for food pellet rewards. During the habituation phase, the maze’s arms were opened, and a 10mg food pellet was baited in each arm. Each mouse was placed in the central zone of the maze and given 10 minutes to explore all arms and consume the pellet. In the four-day training trials, food pellets were baited in arms 1, 6, and 8 with open doors. The remaining arm closed until food pellets were consumed. At this point, they opened for five minutes to let mice enter each arm. For the task test, the arm doors were opened, and food pellets weighing 10 mg were placed in arms 1, 6, and 8. The task ended when the mice consumed food pellets in these arms. The researchers collected data on various parameters, including correct arm entries (CAE), incorrect arm entries (IAE), total arms visited (AV), memory score (MS), working memory errors (WMEs), reference memory errors (RMEs), and total time (TT). Equation 3 was used to calculate the memory score [42].

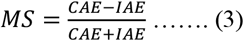

### 2.12 Preparation of brain samples

On day 60, the mice were euthanized using cervical dislocation. Subsequently, the skulls were dissected, and the brains were isolated, meticulously cleansed to eliminate meninges, and rinsed with cold phosphate-buffered saline. Following this, the brains were transferred to microcentrifuge tubes filled with Favorgen™ RNA Stabilization Solution and stored at -20°C until used.

### 2.13 Quantitative polymerase chain reaction (qPCR) for gene expression

Genes expression for neurogenesis (dcx, nestin, Hes5 and NFIA) were analyzed through the qPCR [36]. The mRNA was extracted using the Favorgen™ Tissue Total RNA Mini Kit following the manufacturer’s instructions. The extracted RNA was quantified using a NanoDrop spectrophotometer and then stored at -80°C for further analysis. To synthesize complementary DNA (cDNA), we used the SMOBIO^®^ Reverse Transcription Kit 100RX (cDNA kit) and ten nanograms of isolated RNA as a template. After synthesizing the cDNA, it was stored at -20°C for stability. PCR amplification was performed using the SMOBIO^®^ ExcelTaq 2x PCR Master Dye Mix 100 RXN, with 2.0 μL of DNA template and forward and reverse primers at a concentration of 200.0 nM each. In addition, a total volume of 20.0 μL was achieved by adding 10.0 μL of 2x PCR Master Dye Mix 100 RXN reagents and H2O. The primer sequences for dcx, nestin, Hes5, and NFIA were obtained from the NCBI database, as indicated in Table 1. The specificity of these primers was verified using the Primer Designing Tool (https://www.ncbi.nlm.nih.gov/tools/primer-blast/) and Nucleotide BLAST (https://blast.ncbi.nlm.nih.gov/Blast.cgi?PAGE_TYPE=BlastSearch), both available online at the NCBI website (www.ncbi.nlm.nih.gov). PCR analysis was performed using a T100™ PCR machine. The amplification process included an initial denaturation and enzyme activation step at 95°C for 10 minutes, followed by 40 cycles. Each cycle consisted of denaturation at 95°C for 15 seconds, annealing at 60°C for 30 seconds, and elongation at 72°C for 30 seconds. The final step was elongation at 72°C for 10 minutes. The amplified samples were further analyzed by gel electrophoresis.

**Table 1.**
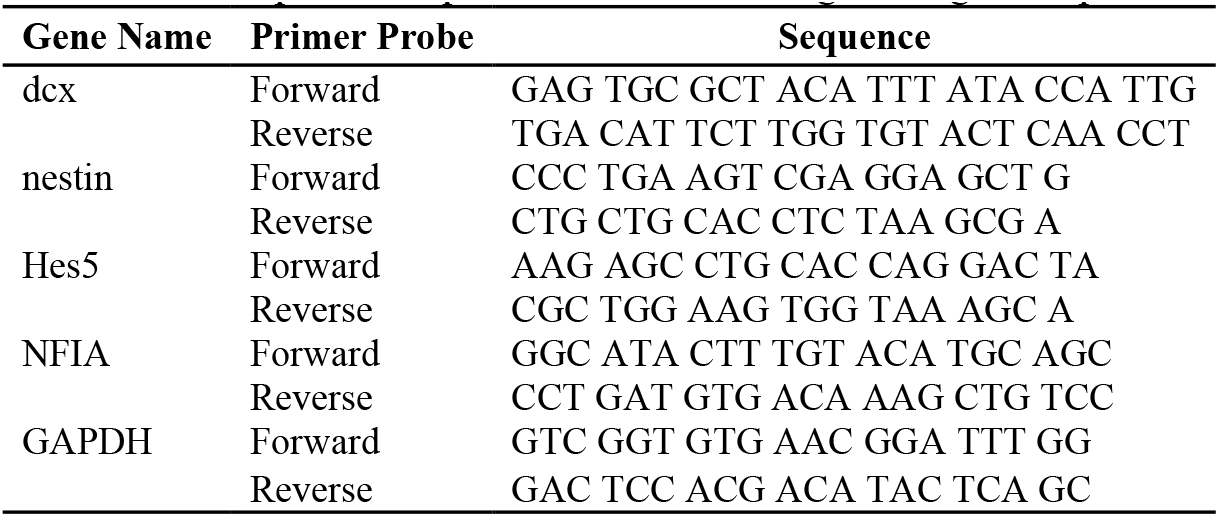
List of primer sequences used for neurogenesis gene expression.

### 2.14 Statistical analysis

The study reports on the results of behavioral assay data (n=10) and gene expression data (n=3), presented as mean ± SEM. The analysis of data normality and homogeneity of variance was conducted using Kolmogorov-Smirnov test and Levene Test. Data differences were evaluated using the Kruskal-Wallis test, supplemented by the Mann-Whitney *U* test, with a significance level set at *P* < 0.05.

## 3. RESULT AND DISCUSSION

### 3.1 Open-field test (OFT)

These studies used the OFT to evaluate locomotor activity and anxiety-like behavior in mice with MSG-induced AD-like symptoms. The findings exhibit no significant differences in travel distance (*P* = 0.442) and grooming time (*P* = 0.307) among all groups (Fig. 1(a) and 1(d)). However, we observed a significant difference in immobility time (*P* = 0.017) and rearing frequency (*P* = 0.005) between HC and AC groups (Fig. 1(b) and 1(c)). Conversely, no significant differences were found in immobility time and rearing frequency when comparing the SP_40, SN_20, SN_40, and DPZ groups to the AC group (*P* > 0.05).

**Figure 1.**
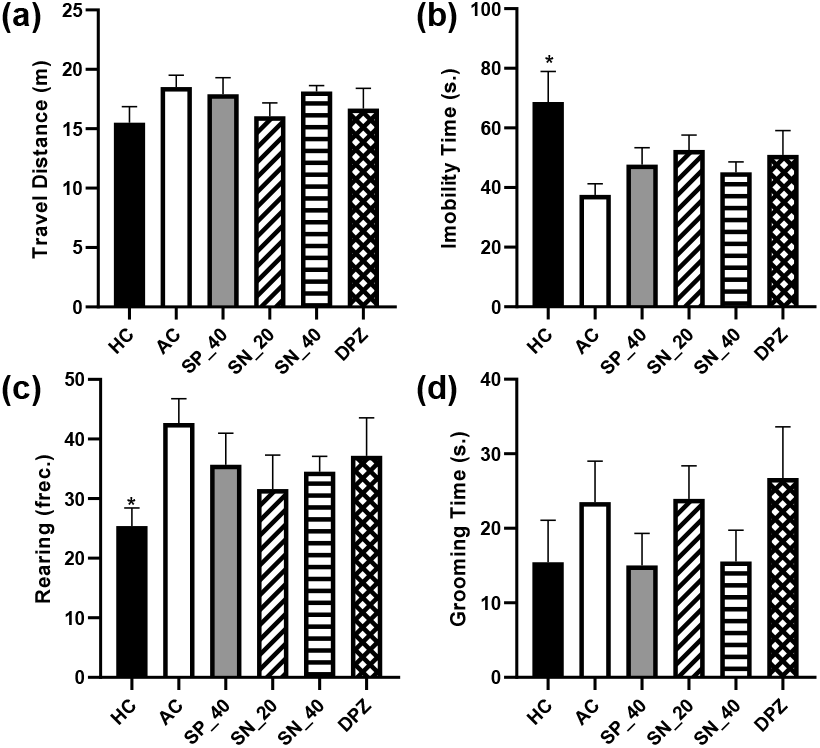
SNEDDS PGV-0 effect on the (a) travel distance, (b) immobility time, (c) rearing and (d) grooming in mice with MSG-induced AD-like symptoms after being treated for six weeks. HC (healthy control), AC (Alzheimer’s control, CMC-Na 0.5%), SP_40 (suspension PGV-0 40 mg/kg), SN_20 (SNEDDS PGV-0 20 mg/kg), SN_40 (SNEDDS PGV-0 40 mg/kg), and DPZ (Donepezil HCl 3 mg/kg). Data presented as mean±SEM (n=10), * *P* < 0.01 vs. AC group by Mann-Whitney *U* test, *ns*. = not significant.

**Figure 2.**
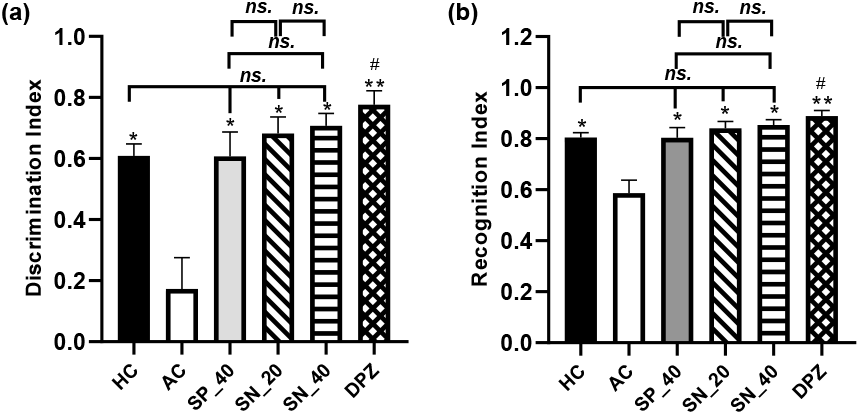
SNEDDS PGV-0 effect on the (a) discrimination and (b) recognition index in mice with MSG-induced AD-like symptoms after being treated for six weeks. HC (healthy control), AC (Alzheimer’s control, CMC-Na 0.5%), SP_40 (suspension PGV-0 40 mg/kg), SN_20 (SNEDDS PGV-0 20 mg/kg), SN_40 (SNEDDS PGV-0 40 mg/kg), and DPZ (Donepezil HCl 3 mg/kg). Data presented as mean±SEM (n=10). *ns*. = not significant (*P* > 0.05), * *P* < 0.01, ** *P* < 0.001 vs. AC group, # *P* < 0.01 vs. HC by Mann-Whitney *U* test.

### 3.2 Novel Object Recognition Task (NORT)

The study found that the AC group exhibited a significant decrease in DI and RI compared to the HC group (*P* = 0.003 and *P* = 0.003, respectively). This result suggests that four-week-old mice treated with MSG experienced a decrease in their ability to recognize new objects. However, there was no significant difference among the SP_40, SN_20, and SN_40 groups compared to the HC group (*P* > 0.05), suggesting that the 45-day treatment of suspension PGV-0 and SNEDDS PGV-0 significantly improved the memory deficit in mice with MSG-induced AD-like symptoms. Moreover, the DPZ group exhibited a significant improvement in recognizing new objects, with higher DI (*P* = 0.008) and RI (*P* = 0.008) than the HC group.

### 3.3 8-Radial arm maze (8-RAM) test

On the 55^th^ day of treatment, which was late in week 7, we conducted a spatial learning and memory assessment using the 8-RAM. The results showed that the AC group had cognitive impairment. There was a significant increase in arm visits (Fig. 3a), total time (Fig. 3c), working memory errors (Fig. 3d), and reference memory errors (Fig. 3e) in the AC group compared to the HC group (*P* = 0.0001, *P* = 0.0002, *P* = 0.0002, and *P* = 0.0001, respectively). As a result, the mice in the AC group had a significant decrease in memory scores (Figure 3b, *P* = 0.0004 vs. the HC group). After a 55-day treatment with suspension PGV-0, both doses of SNEDDS PGV-0 and donepezil-HCl, all spatial learning and memory parameters were normalized (*P* > 0.05 vs. HC group). We found no significant differences in arm visits, memory scores, total time, working memory errors, and reference memory errors between the SP_40, SN_20, SN_40, and DPZ groups and the AC group (*P* > 0.005). However, there were no significant differences observed between the SP_40 vs. SN_40 and SN_20 vs. SN_40 groups in any of these parameters (*P* > 0.05).

**Figure 3.**
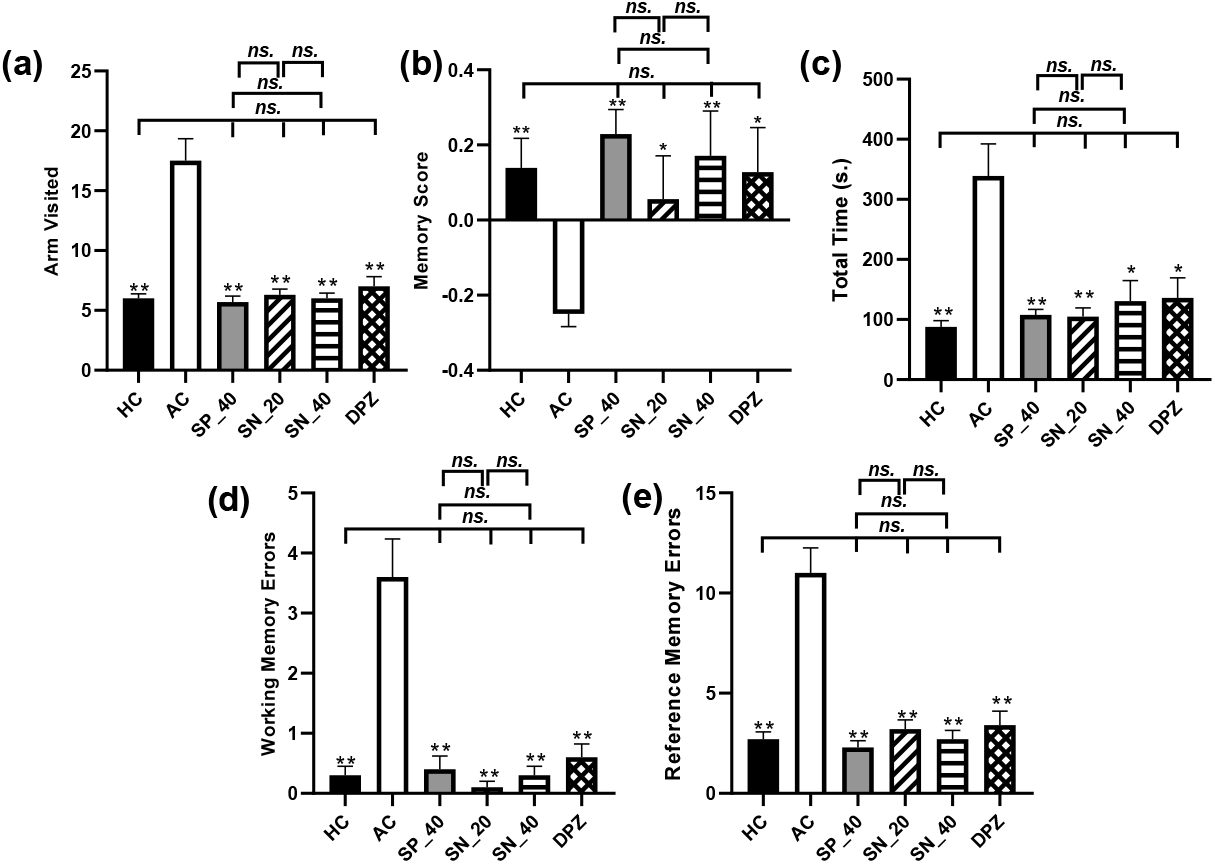
SNEDDS PGV-0 effect on (a) arm visited, (b) memory score, (c) total time, (d) working memory errors, and (e) reference memory errors of mice with MSG-induced AD-like symptoms after seven weeks of treatment. HC (healthy control), AC (Alzheimer’s control, CMC-Na 0.5%), SP_40 (suspension PGV-0 40 mg/kg), SN_20 (SNEDDS PGV-0 20 mg/kg), SN_40 (SNEDDS PGV-0 40 mg/kg), and DPZ (Donepezil HCl 3 mg/kg). Data presented as mean±SEM (n=10). * *P* < 0.01, ** *P* <0.001 vs. AC group by Mann-Whitney *U* test, ns. = not significant (*P* > 0.05)

### 3.4 Effect of SNEDDS PGV-0 on mRNA expression of neurogenesis related genes

The current study examined the impact of SNEDDS PGV-0 on neurogenesis in mice brains exhibiting MSG-induced AD-like symptoms, employing qPCR analysis (Figure 4). Mice in the AC group exhibited a significant decrease in the expression of genes related to neurogenesis, such as dcx, and gliogenesis, such as NFIA (P<0.05). Meanwhile, although not statistically significant, mRNA expression of nestin, a marker of immature neurons, also showed a declining trend in this group. In contrast, the AC group showed elevated mRNA expression of Hes5, suggesting a diminished potential for neurogenesis in the whole brain. However, the treatments involving SNEDDS PGV-0, suspension PGV-0, and donepezil HCl exhibited a significant elevation in the expression of dcx and NFIA and decreasing Hes5 expression (P < 0.01), suggesting amplified neurogenesis in MSG-induced AD-like symptoms in mice. There were no significant differences in the expression of dcx and Hes5 genes between the SP_40, SN_20, SN_40, and DPZ groups and the HC group (P > 0.05). This result implies that treatment with suspension PGV-0, SNEDDS PGV-0, and donepezil-HCl could restore neurogenesis to normal levels in mice with MSG-induced AD-like symptoms.

**Figure 4.**
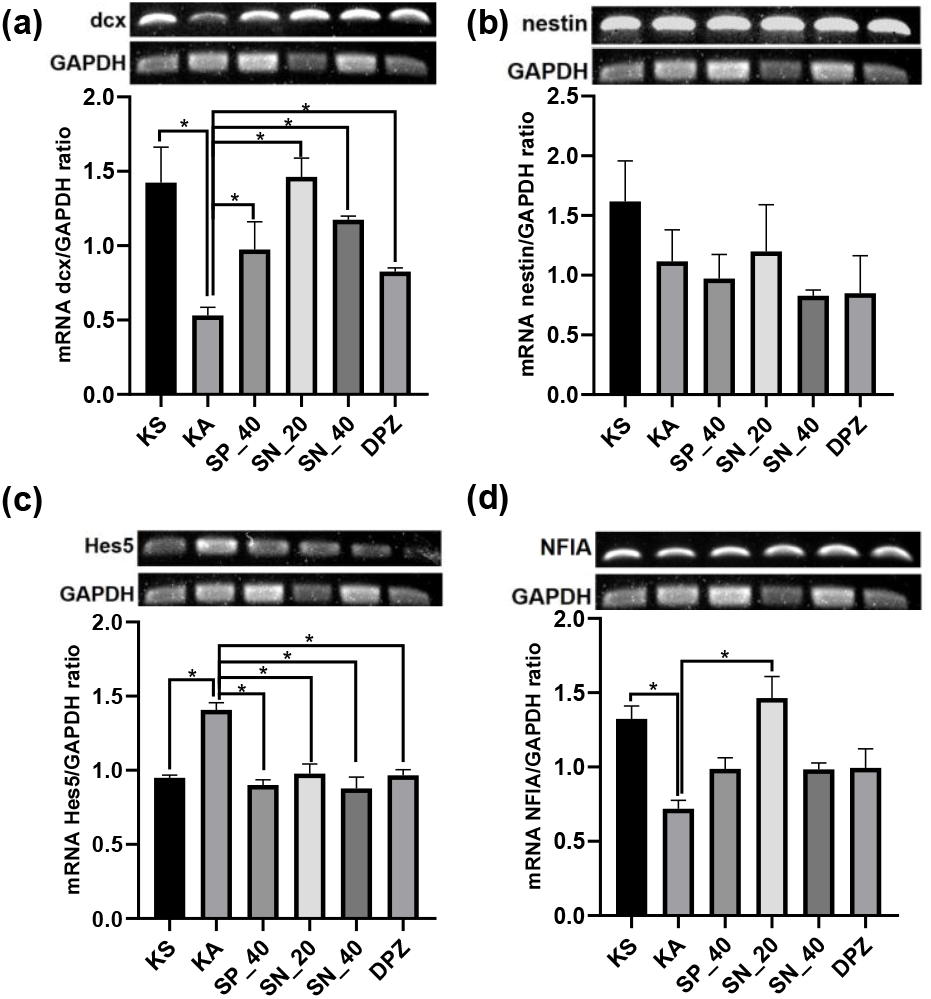
Gene expression ratio of (a) dcx, (b) nestin, (c) Hes5 and (d) NFIA on GAPDH in brain of monosodium glutamate-induced Alzheimer’s mice after treatment for 60 days. Internal GAPDH expression was used to normalize all qPCR data. HC (healthy control), AC (Alzheimer’s control, CMC-Na 0.5%), SP_40 (suspension PGV-0 40 mg/kg), SN_20 (SNEDDS PGV-0 20 mg/kg), SN_40 (SNEDDS PGV-0 40 mg/kg), DPZ (Donepezil HCl 3 mg/kg), dcx (doublecortin), Hes5 (hairy enhancer of split 5) and NFIA (nuclear factor IA). Data presented as mean±SEM (n=3). * *P* < 0.05 vs. AC group by Mann-Whitney *U* test.

### 3.5 Discussion

This study used MSG to induce spatial learning and memory impairment and reduce neurogenesis in Balb/C mice. At four weeks of age, the mice received seven subcutaneous injections of MSG at 4 g/kg. Previous studies have shown that MSG can be toxic and result in memory impairments in animal models. MSG can cause excitotoxicity, leading to significant neuronal damage, particularly in the hippocampus [43–45]. When glutamate accumulates in the synaptic cleft, neurons are overstimulated, leading to apoptosis and neurodegeneration [46,47]. Our recent study showed that when administered to 4-week-old mice, MSG did not affect mice’s travel distance or grooming time. However, it significantly decreased their immobility time and increased rearing frequency (Figure 1). In addition, it reduced mice’s DI and RI on the NORT (Figure 2) and decreased memory score on the 8-RAM task, as shown by the elevated number of AV, WMEs, RMEs, and TT to complete the 8-RAM task (Figure 3).

The findings in our recent study were consistent with earlier studies that suggested MSG exposure may cause long-term damage to the brain, resulting in neurotoxicity, impaired short-term memory, and changes in exploratory behavior [48,49]. Exposure to high levels of MSG in juvenile rodents may induce excitotoxic effects because of their underdeveloped blood-brain barrier [50]. Studies have shown that administering MSG early in life can lead to tau pathology and changes in behavior and cognitive function in rodents [51,52]. Another study found that administering MSG can alter spatial memory and increase Aβ levels [53,54]. Both were hallmark characteristics of AD, especially in the hippocampus [4]. Dissolving MSG in water causes glutamate levels to elevate in the bloodstream, leading to an excessive activation of glutamate receptors. This overstimulation triggers the release of calcium ions, causing damage to vital cellular structures such as the cytoskeleton, cell membrane, and DNA [55]. In addition, MSG can induce oxidative stress, exacerbating its neurotoxicity. It increases malondialdehyde (MDA) levels while decreasing superoxide dismutase (SOD) activity in the brain, both of which are indicators of oxidative stress [44]. The brain is vulnerable to oxidative stress because it has a high metabolic activity and limited antioxidant capacity. Therefore, oxidative stress generates free radicals that can damage cell membranes and DNA, resulting in cell damage and apoptosis [56,57].

Known for its antioxidant [20,21] and anti-inflammatory [17] activities, PGV-0 acts as a potent HDAC2 inhibitor [25]. When administered as a nano-emulsion, PGV-0 inhibits HDAC2 expression [36], an enzyme involved in neurogenesis termination in the mouse brain, linked to memory loss in AD [28]. In the current study, six weeks treatment of PGV-0, either in conventional suspension or loaded SNEDDS, ameliorated cognitive deficits, improved spatial learning, and memory impairment in mice with MSG-induced AD-like symptoms. The mice treated with SNEDDS PGV-0 and PGV-0 suspension showed significant improvement in DI, RI (Figure 2), and MS (Figure 3) compared to the AC group. However, no significant differences in behavioral changes or gene expression related to neurogenesis-gliogenesis were observed in mice with MSG-induced AD-like symptoms following treatment with PGV-0 suspension and both doses of PGV-0 SNEDDS (P > 0.05). The goal of developing PGV-0 into SNEDDS form in this study was to enhance its oral bioavailability. An increase in the oral bioavailability of PGV-0 was observed in the test results following the administration of a single dose of SNEDDS PGV-0 (40 mg/kg.BW), as compared to the PGV-0 suspension at a similar dose. In addition, PGV-0 was detected in brain tissue following PGV-0 suspension treatment, although at lower levels compared to PGV-0 SNEDDS (data not shown and published separately). The lack of difference in effect may be attributed to the timing of testing, which occurred after the administration of PGV-0 suspension and SNEDDS PGV-0 for a period exceeding six weeks. This duration likely allowed for sufficient accumulation of PGV-0 in the brain, leading to alterations in behavior and gene expression related to neurogenesis and gliogenesis. Therefore, further studies are necessary to clarify the difference between administering multiple doses of PGV-0 suspension and SNEDDS PGV-0 in terms of the accumulation of PGV-0 levels in the brain tissue of mice with Alzheimer’s disease.

Neurogenesis is the development of new neurons from neural stem cells. During this process, intermediate progenitor cells differentiate into mature neurons [58]. However, the brains of humans with AD and rodent AD models show abnormalities in neurogenesis [59].

These abnormalities affect different phases of neurogenesis, including the migration, dendritic maturation, and synaptic integration of newborn neurons. Gene expression analysis, employing biomarkers such as the mRNA expression of dcx, nestin, Hes5, and NFIA, facilitates the detection of impaired neurogenesis in the study [58]. The present study exhibited a favorable association between the amelioration of behavioral impairment and altered mRNA expression linked to neurogenesis following a sixty-day administration of SNEDDS PGV-0 in mice with monosodium glutamate-induced Alzheimer’s disease-like symptoms. The crucial role of dcx in neurogenesis lies in its ability to facilitate neuronal differentiation and migration. It also supports stabilizing microtubules, which are essential for neuronal maturation. Its expression is highest in neurogenic zones, such as the subventricular and the sub-granular zone [60]. Our study found that mice treated with PGV-0 suspension, SNEDDS PGV-0, and donepezil-HCl exhibited higher mRNA expression of dcx compared to the AC group. Conversely, the mice treated with PGV-0 suspension, SNEDDS PGV-0, and donepezil-HCl exhibited suppression of Hes5 mRNA expression. A decrease in Hes5 expression promotes pro-neural gene expression and stimulates neurogenesis [61]. Hes5 is a transcriptional repressor with a primary function of repressing pro-neural gene expression, such as Neurog2 and Ascl1, which act as inducers of neuronal differentiation [62]. In addition, SNEDDS PGV-0 (20 mg/kg) enhances mRNA expression of Nuclear Factor IA (NFIA) in mice with monosodium glutamate-induced Alzheimer’s disease-like symptoms. Recent studies have shown the downregulation of NFIA expression around reactive astrocytes in brain tissue from AD patients. Astrocyte reactivity inducing decreased NFIA expression is associated with amyloid-β plaque pathology in brain tissue from AD patients [63]. Previous studies have revealed that the transient upregulation of NFIA can trigger the conversion of neural stem cells into astrocytes and facilitate synaptogenesis [64] to establish connections between new neurons and the neural circuit.

## 4. CONCLUSION

SNEDDS PGV-0 has exhibited efficacy in ameliorating cognitive and memory deficits in mice with MSG-induced AD-like symptoms. The study observed improvements in various behavioral parameters, including DI, RI, and MS. Moreover, the study suggested that SNEDDS PGV-0 restored the expression of critical neurogenesis-related genes, such as dcx and Hes5. The results of this initial study suggest that PGV-0 may promote neurogenesis and could be developed as a drug for Alzheimer’s disease and other neurodegenerative disorders. Further research is necessary to establish the comprehensive effects of PGV-0 SNEDDS as a neurogenesis-promoting agent. Several parameters need monitoring, including the immunostaining results of dcx-positive and nestin-positive cells, NeuN-positive cells (a marker for mature neurons), BrdU-positive cells, GFAP-positive cells, and others in the hippocampus and cortex of Alzheimer’s mice.

## ACKNOWLEDGMENT

The authors wish to express their sincere gratitude to Mr. Panji Pranata, Mr. Surono, and Dr. dr. Nur Arfian for their invaluable technical guidance and support during the study at the Pharmacology Laboratory of the Faculty of Pharmacy and the Anatomy Laboratory of the Faculty of Medicine, Public Health, and Nursing at Universitas Gadjah Mada.

## AUTHOR CONTRIBUTION

The authors confirm their respective roles in this paper as outlined herein. The research idea and study design were developed by Yance Anas and Nunung Yuniarti. Yance Anas conducted the experimental work and collected data, while data analysis, interpretation, and visualization were performed by Yance Anas and Ronny Martien. Ratna Asmah Susidarti and Nunung Yuniarti proofread the data. Yance Anas and Nunung Yuniarti collaborated in drafting the manuscript. After a thorough review of the results, all authors agreed to the final manuscript.

## FINANCIAL SUPPORT

This work was funded by the Final Project Recognition Grant from Universitas Gadjah Mada (Grant Number 5075/UN1. P. II/Dit-Lit/PT.01.01/2023).

## CONFLICT OF INTEREST

The authors assert that there are no conflicts of interest related to the publication of this original article.

## ETHICAL APPROVALS

All study procedures were approved by the ethics commission of Integrated Testing and Research Laboratory Universitas Gadjah Mada, Indonesia, under license number 00016/04/LPPT/VI/2022.

## DATA AVAILABILITY

The first author holds the primary and secondary data referenced in this manuscript, and agrees to make it available on demand.

## PUBLISHER NOTE’S

## Notes

### Competing Interest Statement

The authors have declared no competing interest.

